# mixKIN: An ancestry-aware kinship estimation method for low coverage genomes

**DOI:** 10.64898/2026.10.13.758828

**Authors:** Muhammed Sıddık Kılıç, Anna-Sapfo Malaspinas, Dilek Koptekin

## Abstract

Genetic kinship analysis provides insights into the social and biological organization of past populations. However, kinship estimation from low-coverage genomes, which are common in ancient DNA (aDNA) studies, remains challenging, particularly in genetically heterogeneous or admixed populations. Approaches that assume a homogeneous background relatedness can bias estimates when individuals have distinct ancestry. We developed ***mixKIN***, an ancestry-aware kinship estimation from low-coverage genomes. Rather than using a single mismatch baseline for all pairs, *mixKIN* estimates pair-specific baseline by weighting within-population diversity and between-population divergence according to ancestry proportions of both individuals in each pair. We compared *mixKIN* with *correctKin* and *RelateAdmix*, which account for population structure and admixture, and with *READv2* representing a widely used approach that assumes a homogeneous population background. To benchmark the methods, we ran extensive pedigree simulations across diverse demographic scenarios, including distant-population admixture, intra-population heterogeneity, and genetic outliers, at coverages from 1x to 0.03x. *mixKIN* achieved higher average classification performance in admixed and heterogeneous scenarios, while maintaining comparable performance in homogeneous populations, with relationships evaluated up to third-degree. In an application to an empirical data from a medieval English cemetery, *mixKIN* recovered relationships involving an individual with West African and European ancestries without the need for a manual adjustment to account for admixture. These findings support ancestry-aware mismatch normalization as a practical approach to improving kinship estimation and retaining genetically diverse individuals in kinship studies.

## Background

Estimating genetic kinship from ancient genomes contributes to reconstructing family relationships and understanding past social organization. Reconstructed pedigrees, interpreted alongside archaeological evidence, can inform our understanding of how communities were organized, how resources were shared, and how individuals moved between groups. Several ancient European settlements have shown patterns consistent with patrilocal residence (Fowler et al., 2022; Gelabert et al., 2025; Gnecchi-Ruscone et al., 2024; Mittnik et al., 2019; Rivollat et al., 2023; Szécsényi-Nagy et al., 2015), while other studies have identified female-centered and/or matrilocal residence such as in Çatalhöyük in Neolithic Central Anatolia (Yüncü et al., 2025), Fujia in Neolithic China (Wang et al., 2025), and Durotrigian Iron Age Britain (Cassidy et al., 2025). Kinship analyses have also provided insight into social stratification and marriage strategies, including polygamous unions (Blöcher et al., 2023) and levirate customs (Gnecchi-Ruscone et al., 2024), while helping address a wide range of other anthropological, social, and genetic questions.

Despite these important biological and cultural insights, reconstructing genetic relationships - which relies on kinship estimation - is a difficult task in ancient DNA. In particular, estimating kinship is very challenging for low coverage genomes, which account for a large portion of the data in ancient DNA studies. Moreover, population structure and admixture further complicate kinship estimation, particularly when individuals differ substantially in their genetic profile. Haplotype-based approaches can reduce sensitivity to population structure but generally require high-coverage genomes or genotype imputation (Ringbauer et al., 2024); however, reliable imputation remains difficult for many ancient samples, especially for genomes with a depth of coverage below 0.5x (Sousa da Mota et al., 2023). There exist several approaches for low-coverage data such as *NgsRelate* (Korneliussen & Moltke, 2015), *READ/READv2* (Alaçamlı et al., 2024; Monroy Kuhn et al., 2018) and *KIN* (Popli et al., 2023). In these methods, a background relatedness is being computed to get a sense of the expected level of genetic similarity between unrelated individuals in the population. Yet, all these methods typically assume a homogeneous background across the individuals being analysed, an assumption that can lead to biased estimates in genetically heterogeneous or admixed populations. In particular, normalization against a shared population baseline - the expected level of genetic similarity between unrelated individuals in the population - can underestimate or overestimate kinship between individuals with variable admixture proportions (Ringbauer et al., 2024; Sayer et al., 2025). Such biases can lead to missed relationships involving individuals with distinct ancestry profiles and may prompt their exclusion from kinship analysis. Methods such as *correctKin* (Nyerki et al., 2023) and *RelateAdmix* (Moltke & Albrechtsen, 2014) explicitly address population structure or admixture, but their performance has been shown to be limited (Lefeuvre et al., 2026).

We therefore developed *mixKIN* to account for ancestry differences when estimating genetic kinship. Rather than assuming a homogeneous background relatedness, *mixKIN* estimates a baseline for each pair by weighting within-population diversity and between-population divergence accounting to the individual’s ancestry proportions (see Methods). We evaluated *mixKIN* performance using simulated pedigrees spanning homogeneous, heterogeneous, and recently admixed populations, including datasets with admixed outliers. We compare our approach with *correctKin*, *RelateAdmix*, *READv2*. Finally, we also applied *mixKIN* to a previously studied family from a medieval English cemetery. Overall, our simulations and empirical application demonstrate that *mixKIN* provides an ancestry-aware framework that improves kinship estimation in admixed and genetically heterogeneous ancient populations compared to all tested methods for depth of coverage ranging from 1x to 0.03x and with relationships evaluated up to the third degree. Notably, *mixKIN* also maintains comparable performance in homogeneous populations. Finally, our application demonstrates the advantage of our approach for real data; we show it is possible to recover the expected relationships by using *mixKIN* and without having to manually correct estimates to account for admixture as was done in the original publication (Sayer et al., 2025).

## Results

### Benchmarking across demographic scenarios

To evaluate our approach, ***mixKIN***, we compared it with *correctKin* (Nyerki et al., 2023) and *RelateAdmix* (Moltke & Albrechtsen, 2014) - which account for population structure and admixture, and with *READv2* (Alaçamlı et al., 2024) the common approach in aDNA to estimate kinship from low coverage genomes.

We considered several demographic scenarios and simulated seven independent pedigrees per scenario. Each pedigree in our simulation includes 14 individuals with up to third degree relationships (see Figure 1A). We combined the resulting 98 (7×14) individuals for kinship estimation. Founders were drawn from either one population or multiple populations, depending on the demographic scenario (Figure 1A). These scenarios included (i) <u>Distant-population admixture:</u> a dataset with admixed Japanese (JPT) and Toscani (TSI) individuals in the pedigrees; (ii) <u>Genetic outliers</u>: a predominantly JPT dataset comprising only JPT founders from six JPT-TSI pedigrees and one complete JPT-TSI pedigree, in which TSI founders and admixed individuals would be considered genetic outliers because their genetic profile differs from that of the predominantly JPT population; (iii) <u>Intra-population heterogeneity</u>: a Peruvian (PEL) population which has high intra-population heterogeneity including individuals with various levels of European ancestry; (iv) <u>Close-population admixture</u>: a dataset with admixed individuals from Toscani (TSI) and Utah Residents with Northern and Western European Ancestry (CEU) populations, (v) <u>Homogeneous population</u>: a homogeneous Japanese (JPT) population. Overall, the pedigrees capture various familial relationships (e.g., siblings, parent-offspring, first cousins) and were evaluated at coverages ranging from 1x to 0.03x (see Methods). Figure 1B summarizes the ancestry-aware pairwise mismatch ratio modelling used by *mixKIN*.

**Figure 1:**
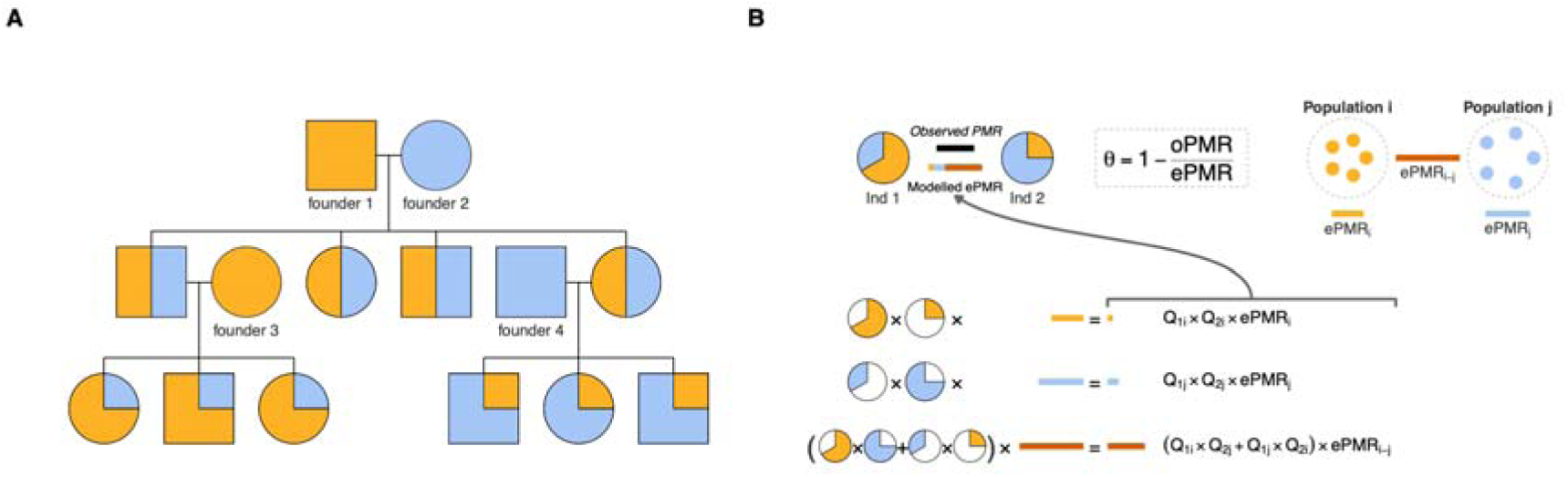
Pedigree simulation structure and the *mixKIN* framework. **(A)** Pedigree structure used in the simulations. Nodes are colored according to the individuals’ ancestry proportions. **(B)** Conceptual overview of the *mixKIN* algorithm for a two-way admixture model. Colored solid lines represent expected Pairwise Mismatch Ratio (ePMR). Intra-population diversities (*ePMR_i_* and *ePMR_j_*) and inter-population divergence (*ePMR_i_*_−*j*_) are derived from the allele frequencies of the source-population (population *i* and *j*). For each pair, ePMR calculated as the sum of intra- and inter-population ePMRs, adjusted by ancestry proportion (*Q*) combinatorial weights. The estimated kinship coefficient (θ) is θ=1-(oPMR/ePMR), where oPMR is the observed pairwise mismatch ratio, shown as solid black line. As illustrated in the example, an oPMR lower than the modelled ePMR indicates genetic kinship between the two individuals.

We then used several approaches to estimate ancestry proportions for the methods accounting for population structure. Individuals’ ancestry proportions were primarily estimated using qpAdm (Haak et al., 2015) for *mixKIN* and unsupervised *ADMIXTURE* (Alexander et al., 2009) for *RelateAdmix*. In the distant-admixture scenarios, both methods were also tested by using supervised *ADMIXTURE* to estimate ancestry proportion. Finally, kinship estimates were adjusted after *PCAngsd* analysis with external reference panels for *correctKin* (see Methods).

We assessed relationship classification using one minus Uniform Ordinal Classification (1-UOC) index, an ordinal performance metric that penalizes errors according to the distance between true and predicted relationship categories. We additionally evaluated kinship estimates using root mean square deviation (RMSD) and mean bias error (MBE), which quantify error magnitude and direction, respectively (see Methods).

### Kinship estimation under distant-population admixture

In the distant-population admixture scenario (JPT-TSI), *mixKIN* achieved the highest classification scores across the evaluated coverage range, maintaining 1-UOC values above 0.93 for depth of coverages from 1x to 0.1x. None of the methods that we compared to *mixKIN* reached a higher 1-UOC index level even at 1x coverage (Figure 2A and 2B).

**Figure 2:**
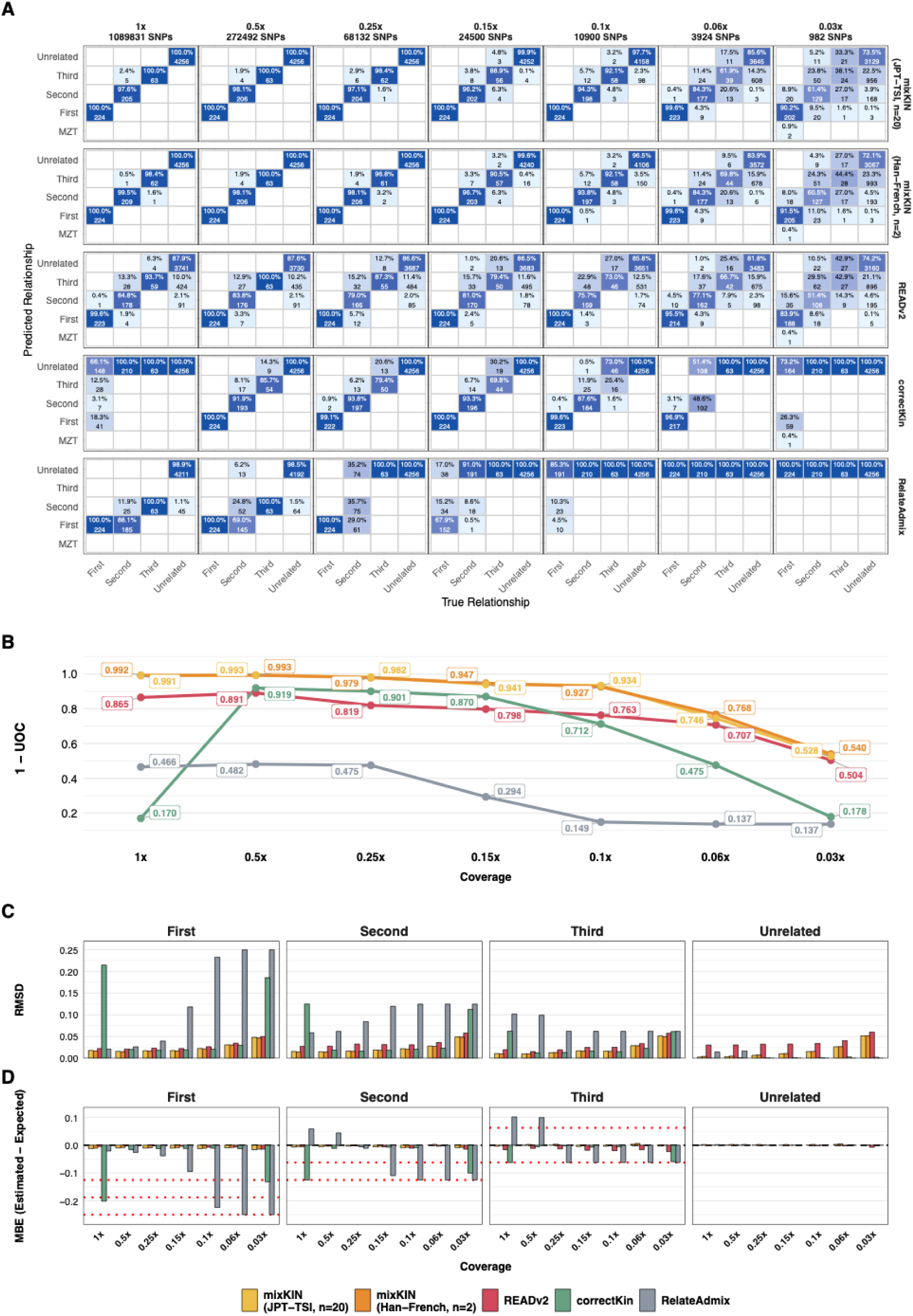
Comparison of kinship estimation methods in the distant-population admixture scenario (JPT-TSI). **(A)** Confusion matrices comparing true and predicted relationships for each method. Each cell shows the number and percentage of pairs assigned to corresponding predicted relationship categories. Text within each cell is written in black for percentages below 50% and in white for percentages of 50% or higher. Diagonal cells represent correct classifications. Darker blue shading indicates higher values across all cells. The average number of overlapping SNPs is shown for each coverage level on top of each column. **(B)** Classification performance measured using the 1-UOC index across 1x to 0.03x coverage, higher values indicate better performance. *mixKIN* relying on true source populations (yellow) or close-proxy populations (orange) is compared with *correctKin*, *RelateAdmix* and *READv2*. **(C)** Root Mean Square Deviation (RMSD) between estimated and expected kinship coefficients, stratified by true relationship degree. **(D)** Mean Bias Error (MBE) calculated as *θ_est_*-*θ_exp_*, within each true relationship category. Positive and negative values indicate overestimation and underestimation, respectively. Red dotted lines indicate deviations corresponding to one degree of relatedness in either direction.

Among the other methods accounting for population structure, *correctKin* performed better at intermediate coverages (0.5x-0.1x), but its performance declined after 0.1x. Interestingly, at 1x coverage, its kinship estimates were unchanged from the *PCAngsd* output, indicating no effective correction was applied; the reason for this behavior is unclear. *RelateAdmix* showed the lowest overall classification performance. On average, READv*2* performed worse than *mixKIN* but it achieved better average performance across all evaluated coverages compared to all other tools (mean 1-UOC: 0.764; range: 0.891 - 0.504). Interestingly, *READv2*’s performance was slightly lower than *correctKin* over the intermediate coverage.

Comparisons of estimated kinship coefficients within each relationship category showed that *mixKIN* achieved the lowest average RMSD across coverage levels for first-, second-, and third-degree relationships (Figure 2C). For unrelated pairs, *correctKin* and *RelateAdmix* had lower reported RMSDs than *mixKIN*, but these comparisons were affected by differences in output handling: *correctKin* estimates are unavailable for pairs classified as unrelated and assigned zero, whereas *RelateAdmix* estimates are constrained to be non-negative. These ad-hoc thresholds and values limit direct comparisons of RMSD and MBE for unrelated pairs. Nevertheless, *mixKIN* showed a lower average RMSD than *READv2* in this category.

*mixKIN* had the smallest average absolute MBE for second- and third-degree relationships across methods. *READv2* performed slightly better than *mixKIN* for first-degree relationships but it performed slightly worse for other relationships. MBE further revealed *correctKin* is systematically underestimating for first-, second-, and third-degree relatives. Moreover, *RelateAdmix* is overestimating second-, and third-degree relatives for 1x and 0.5x, and is always underestimating relatedness with lower coverage levels (< 0.5x) (Figure 2D). We also note that, for unrelated pairs, all methods had average MBE values close to zero, although these values should be interpreted in light of the output-handling differences described above.

#### Using close-proxy populations

We also run *mixKIN* by using only two diploid individuals from Han and French instead of twenty diploid individuals from JPT and TSI to estimate individuals’ ancestry proportions and allele-frequencies. *mixKIN*’s performance was almost identical. These results show that close-proxy populations can be used when the exact source populations are unavailable or represented by very few individuals (Figure 2). *mixKIN* could therefore estimate expected Pairwise Mismatch Ratio (ePMR) using very small close-proxy panels, provided that a large number of loci are available, as inter-locus variance generally exceeds intra-locus variance (Nei & Roychoudhury, 1974).

For the same distant-population admixture scenario, *mixKIN* maintained similar classification performance when using ancestry proportions estimated by supervised *ADMIXTURE* instead of *qpAdm*. In these cases, 1-UOC scores of at least 0.9 down to 0.1x coverage were observed (Figure 3; Supplementary Figure 1). Note that with supervised *ADMIXTURE* inputs, *RelateAdmix* showed substantially lower performance (mean 1-UOC: 0.454, range: 0.478 - 0.413). Although it correctly identified all first-degree pairs, it classified more than 85% of second-degree pairs as first-degree, averaged across coverage levels, indicating overestimation of relatedness.

**Figure 3:**
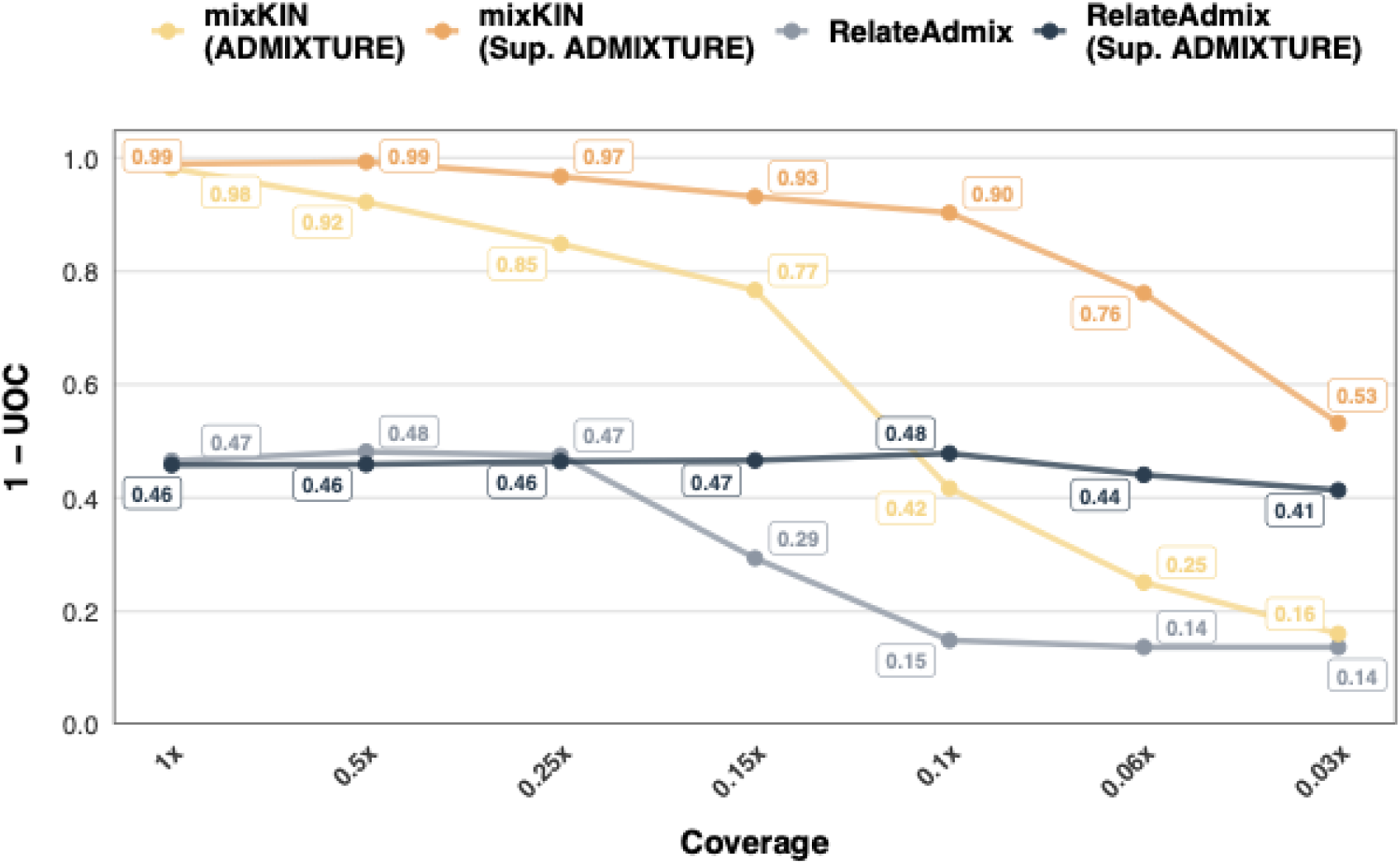
Performance comparison of *mixKIN* and *RelateAdmix* when ancestry proportions estimated using *ADMIXTURE* in the distant-population admixture scenario (JPT-TSI). Individual ancestry proportions and source population allele-frequencies were estimated using either unsupervised *ADMIXTURE* or supervised *ADMIXTURE* (Sup. ADMIXTURE). Performance is measured using the 1-UOC index across coverages (1x to 0.03x).

With unsupervised *ADMIXTURE* inputs, *RelateAdmix* performed similarly to its supervised *ADMIXTURE* analysis down to 0.25x coverage (1-UOC: ∼ 0.47), but performance declined sharply below this level, with most pairs classified as unrelated (Figure 3; Supplementary Figure 1). *mixKIN* also showed a marked decline at coverages below 0.1x (mean 1-UOC across all evaluated coverages: 0.621; max: 0.981; min: 0.160). The loss of accuracy at low-coverage of both methods is consistent with less reliable ancestry proportions and allele-frequency estimates at low-coverage.

### Recovering kinship for a population including genetic outliers

To evaluate a setting commonly encountered in empirical aDNA datasets, a largely homogeneous population containing a few individuals with distinct ancestry, we combined only JPT founders from six JPT-TSI pedigrees with one full JPT-TSI pedigree. Here, TSI founders and admixed individuals from JPT-TSI pedigree represented genetic outliers.

As above, *mixKIN* is outperforming the other methods. Furthermore, *mixKIN* maintained its classification performance despite ancestry imbalance, recovering relationship degrees involving these individuals thanks to its pair-specific ancestry-aware baselines (Figure 4A; Supplementary Figure 2).

**Figure 4:**
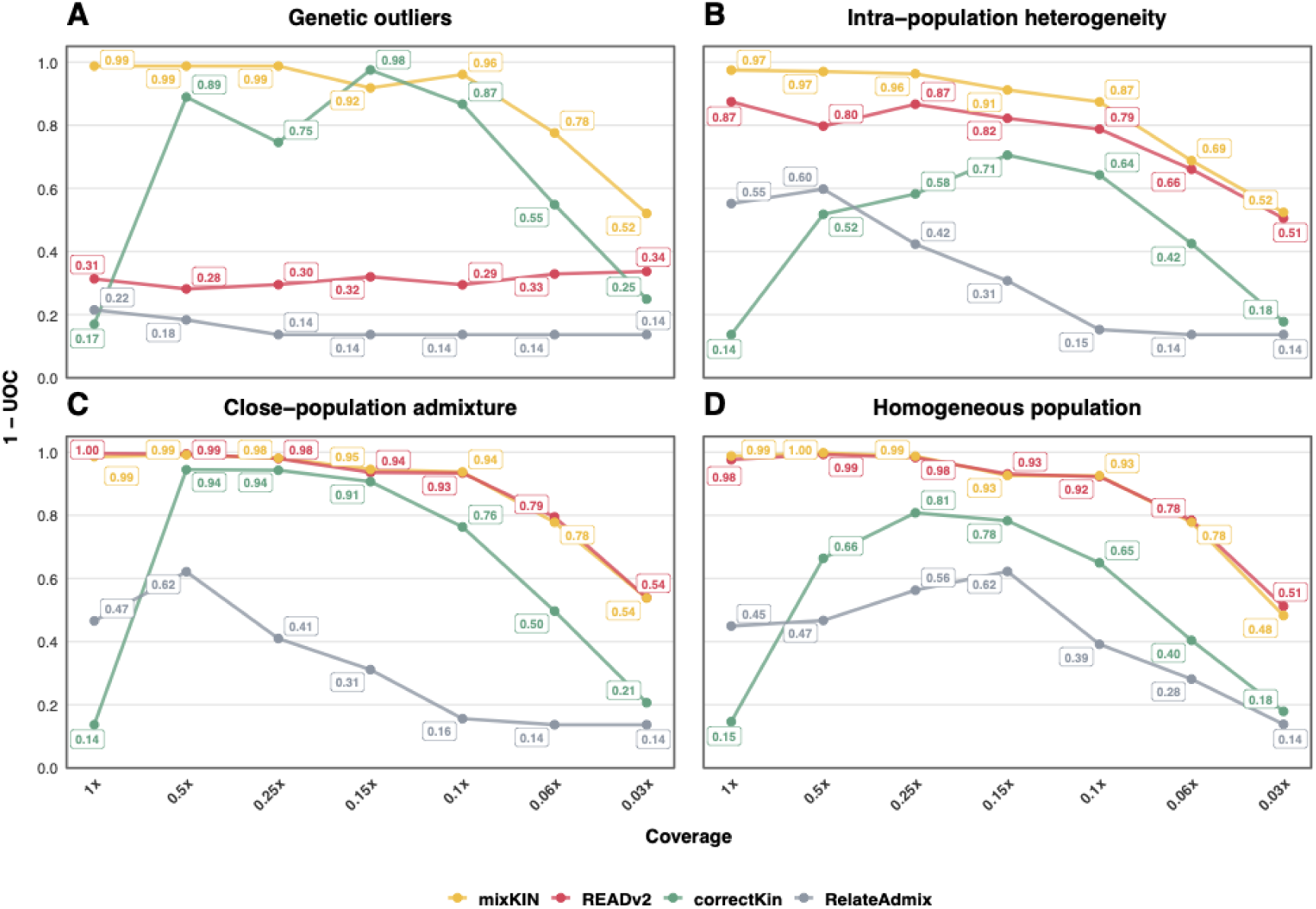
Classification performance across demographic scenarios. Performance is measured using the 1-UOC index across coverages from 1x to 0.03x. Panels show: **(A)** Genetic outliers, **(B)** intra-population heterogeneity within Peruvian population, **(C)** close population admixture between TSI and CEU, and **(D)** a homogeneous JPT population. Reference sample sizes used by *mixKIN* for allele frequency estimation were Mayan (*n* = 18) and IBS (*n* = 103) for the PEL, *n* = 2 per source population for TSI–CEU, and *n* = 2 for JPT.

*correctKin* showed a similar coverage-dependent pattern to the balanced distant-admixture scenario: performance was stronger between 0.5x and 0.1x, whereas estimates at 1x remained not corrected. *RelateAdmix* had the lowest classification scores 1-UOC with values approximately 0.12 across different coverages, potentially reflecting difficulties in estimating ancestry proportions with unsupervised *ADMIXTURE* when ancestral populations are unevenly represented (Shringarpure & Xing, 2014).

*READv2* also showed noticeably reduced performance (1-UOC ∼ 0.3), underestimating kinship for admixed pairs and frequently assigning them to more distant relationship categories (Figure 4A; Supplementary Figure 2A). This might be caused by background relatedness estimated from the median of PMRs in *READv2*, which does not reflect well the actual background relatedness of pairs involving individuals with TSI ancestry.

### Kinship estimation within a single population with high intra-population heterogeneity

We evaluate kinship estimation within a single heterogeneous population in which individuals have varying ancestry proportions of distant source populations. For this scenario, we used the Peruvian (PEL) population, which exhibits within-population variation in European and Indigenous ancestry proportions. We selected founders whose European-related ancestry ranged from 2-37% as estimated with *qpAdm* using Iberian (IBS) and Mayan as source populations.

As for the other scenarios, *mixKIN* achieved the highest average classification performance among the evaluated methods (Figure 4B; Supplementary Figure 3). *READv2* - which does not account for admixture - came second, with better overall 1-UOC values than *correctKin* and *RelateAdmix* (mean 1-UOC: 0.759; range: 0.875 - 0.505). Among the other ancestry-aware approaches, *correctKin* performed better than *RelateAdmix*, but both had considerably lower average scores (mean 1-UOC: 0.455 and 0.329 respectively) than *mixKIN*. Note that as in the distant-admixture scenario, unexpectedly, *correctKin* showed no correction at 1x coverage leading to a very low 1-UOC value. These results highlight the importance of ancestry-aware, pair-specific baselines not only in populations admixed within past few generations, as in JPT-TSI scenario, but also within a single population where variation in individual ancestry proportions can bias kinship estimation assuming a uniform, homogeneous population background.

### Kinship estimation in homogeneous and closely related admixed populations

In the close-admixture scenario involving the two European, TSI and CEU populations, *mixKIN* achieved higher average classification performance than *correctKin* and *RelateAdmix* (Figure 4C; Supplementary Figure 4). These three methods showed broadly similar performance to that observed in the distant admixture scenario, with mean 1-UOC scores of 0.880, 0.628 and 0.320, respectively. In contrast, *READv2* improved from a mean 1-UOC of 0.764 in the distant-admixture scenario to 0.882, achieving performance slightly higher than *mixKIN*. The improvement in *READv2* is consistent with the limited genetic differentiation between TSI (within-population ePMR: 0.2701) and CEU (within-population ePMR: 0.2696), such that the combined dataset can be treated as approximately homogeneous for PMR normalization.

In the homogeneous population scenario with the Japanese (JPT) founders, we applied *mixKIN* without ancestry modelling, estimating ePMR directly from source-population allele frequencies. *mixKIN* again achieved higher average *classification* scores than *correctKin* and *RelateAdmix* and performed similarly to *READv2* (mean 1-UOC: 0.870 and 0.872, respectively) (Figure 4D; Supplementary Figure 5).

### Empirical case: Recovering family relationships at Eastry Updown, a medieval England cemetery

Finally, we applied *mixKIN* to an empirical case from the Eastry Updown cemetery in medieval England. Sayer et al. (2025) have shown that the individuals buried at this cemetery had mixed European and African ancestry.

We focused on a genetic outlier with ∼30% West African ancestry (EAS003; Figure 5), whose relatives were identified as EAS001, EAS002, and EAS006 by manually replacing the population-average baseline with an individual-specific baseline (Sayer et al., 2025).

**Figure 5:**
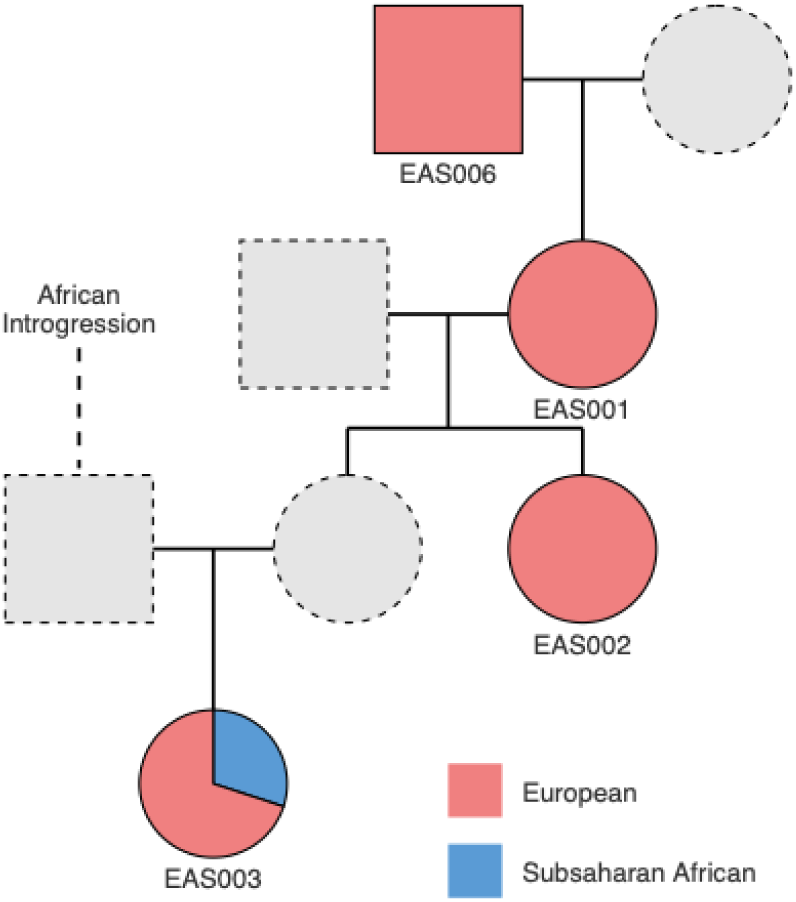
Estimating kinship in an admixed pedigree from the medieval Eastry Updown cemetery. Sampled individuals (solid outlines) and unsampled relatives (dashed outlines) are shown, with node colors indicating European (red) and Sub-Saharan African (blue) ancestry. Adapted from Sayer et al. (2025).

In particular, Sayer et al found that uniform population baseline of 0.255, yields kinship coefficients (*θ*) of 0.0215, 0.0266, and -0.0127 for EAS003 paired with EAS001, EAS002, and EAS006, respectively. In other words, when one does not account for admixture, all three pairs are classified as unrelated (see Supplementary Note 3, Supplementary Table S11). Accounting admixture affects the kinship estimation, Sayer et al. (2025) manually replaced the baseline of 0.255 by an individual-specific baseline 0.275. This allowed them to conclude that EAS003 has three relatives.

We ran *mixKIN* for all individuals in the pedigree (Figure 5) and estimated kinship coefficients (*θ*) of 0.101, 0.106, and 0.067 for the same pairs corresponding to second-, second, and third-degree relationships with EAS003 (see Methods). These classifications are in line with the relationships reported by Sayer et al. (2025).

## Discussion

Our results showed that accounting for ancestry differences improves kinship estimation in genetically admixed ancient populations. Importantly, our approach also performs similar or better than other methods for homogeneous populations. However, when assuming a homogeneous background, the median pairwise mismatch ratio (PMR) primarily reflects the majority population. This can lead to underestimation of kinship involving genetic outliers/admixed individuals. *mixKIN* overcomes this challenge by accounting for these differences by weighting within-population diversity and between-population divergence according to each pair’s ancestry proportions.

The performance improvement is especially evident in the distant-population admixture and genetic outlier scenarios, where we find *mixKIN* having a higher 1-UOC, having more accurate and less biased results than all other methods for almost all tested scenarios from 1x to 0.03x coverages.

The intra-population heterogeneity scenario also shows that current methods also underperform even when admixture started earlier than a few generations, as in the distant-population admixture scenario. Even within a single population, variation in ancestry proportions among individuals can make the assumption of a homogeneous population background unsuitable for kinship estimation.

When considering close-population and or a single homogenous population, *READv2*, which does not account for population structure, and *mixKIN* perform similarly. In those cases, a homogeneous baseline seems to be a reasonable approximation.

An important observation is that our method also performs well under challenging conditions. The close-proxy scenario suggests that neither large reference panels nor exact source populations are always necessary for baseline PMR estimation. Our *mixKIN* results are almost identical when 20 diploid individuals from each exact source population are used, or when only two diploid individuals from close-proxy populations are used as source, suggesting the method works well if enough SNPs are available.

The benchmark results also reveal that accounting for population structure does not always remove all sources of estimation error. For instance, *correctKin* adjusts PCAngsd-derived kinship estimates for SNP overlap, but its accuracy still varies across coverage levels. Its unchanged estimates at 1x coverage require further investigation. *RelateAdmix* explicitly accounts for admixture, yet its performance depended on the ancestry-estimation configuration. Both *RelateAdmix* and *mixKIN* lost accuracy with unsupervised ADMIXTURE-derived inputs at low coverage consistent with less reliable ancestry proportions and source allele frequencies (Alexander et al., 2009; Shringarpure & Xing, 2014).

The Eastry Updown case illustrates how a shared baseline can obscure family relationships involving even one admixed individual. The relatives of the admixed individual (EAS003) could have remained undetected if her distinct genetic profile had not prompted a reassessment of the mismatch baseline. Missing these relationships could have altered interpretations of mobility and integration within the community. *mixKIN* recovered the relationship reported by Sayer et al. (2025) without the manual baseline adjustment used in the study. Such adjustments become very impractical and error prone when several individuals have distinct genetic profiles, because a baseline suitable for one pair may not suit another. We argue that ancestry-aware normalisation provides a systematic alternative to selecting these values individually.

## Limitations and future directions

Although *mixKIN* performed well across the tested demographic scenarios, its application involves a few key methodological considerations. *mixKIN*’s ancestry-aware estimation requires pre-estimated individual admixture proportions and source-population allele frequencies, and its accuracy depends on the quality of these upstream inputs. Additionally, the current framework does not account for inbreeding, which can inflate estimated kinship coefficients, requiring careful interpretation in highly consanguineous populations.

## Conclusions

*mixKIN* provides an ancestry-aware framework for kinship estimation for low-coverage ancient genomes by replacing a shared mismatch baseline with pair-specific expectations. It improved classification in mixed and heterogeneous populations while maintaining comparable performance in homogeneous populations. These findings support ancestry-aware, pair-specific baselines as a practical approach to kinship estimation for cases with and without population structure, helping retain genetically diverse individuals who might otherwise be excluded because a shared population baseline is unsuitable.

## Methods

### mixKIN Algorithm

#### Mismatch-based kinship estimation

*mixKIN* estimates kinship coefficient *theta* (*θ*) between two target individuals. It uses as input pairwise genetic data for those individuals, as well as admixture proportions and allele frequencies of the source populations. We first describe the case where none of the target individuals and then expand our model to the case with admixture.

*mixKIN* estimates kinship coefficient *theta* (*θ*) for each pair of individuals by comparing their observed pairwise mismatch ratio (oPMR) with the expected pairwise mismatch ratio for unrelated individuals (ePMR) over a set of M genetic loci, typically assumed to be distributed genome-wide. Then kinship coefficient is then calculated as:

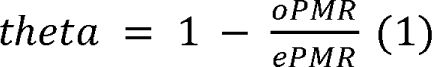

Here, oPMR is the proportion of non-matching genotype calls between two individuals at overlapping sites.

*mixKIN* estimates ePMR directly from source-population allele frequencies. For a single homogeneous population, ePMR represents the probability that two independently sampled alleles differ, averaged across all SNPs used:

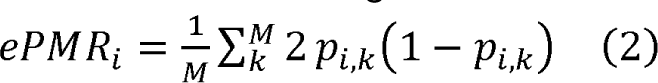

where *p_i,k_*is the allele frequencies at the *k*-th site for the populations *i*, and *M* is the total number of sites.

The ePMR within the population corresponds to the expected average heterozygosity. Because allele-frequencies estimated from a finite number of samples produce a downward-bias in the estimated diversity, we apply the finite-sample correction described by Nei (1978). Since the exact correction depends on the number of observed SNPs at each locus, our implementation approximates the correction factor as 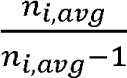. Here, *n_i_*_,*avg*_ is the average number of successfully called alleles per locus in source population *i*. We calculate *n_i_*_,*avg*_ by multiplying the total number of individuals by the average genome-wide genotype call rate and their ploidy (1 for pseudohaploid, 2 for diploid data). The unbiased ePMR is then:

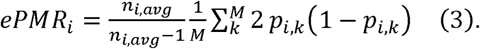

#### Ancestry-aware mismatch baseline

When a dataset contains admixed individuals or individuals with different ancestry profiles, a baseline computed based on a single homogeneous population may not accurately represent every pair. The expected mismatch depends not only on diversity within each ancestral source but also on divergence between sources. *mixKIN* accounts for these differences by combining source-population allele frequencies with the ancestry proportions of both individuals. In the current implementation, individual ancestry proportions and source allele frequencies are assumed to be known. Those can be estimated using methods such as *qpAdm* (Haak et al., 2015) or *ADMIXTURE* (Alexander et al., 2009) before applying *mixKIN*.

For a two-source admixture model, the pair-specific baseline contains three components. The first represents the expected mismatch when both sampled alleles originated from the first source. The second represents the expected mismatch when both sampled alleles originated from the second source. The third represents the expected mismatch when the alleles originate from different sources. For example, consider two unrelated individuals, each with 50% ancestry from source population *i* and 50% from source population *j*. Under independent ancestry draws, there is a 25% probability that both samples’ alleles originated from *i*, a 25% probability that both originate from *j*, and a 50% probability that they originate from different sources. Their expected mismatch baseline therefore combines 25% of the within-*i* mismatch, 25% of the within-*j* mismatch, 50% of the between-source mismatch.

In *mixKIN*, these three components are implemented by weighting the mismatch expectation for source population *i* and *j* as follow:

i. The probability that both alleles are drawn from source population *i*, multiplied by the within population ePMR for *i* (corrected heterozygosity for population *i*).
ii. The probability that both alleles are drawn from source population *j*, multiplied by the within-population ePMR for *j*.
iii. The probability that alleles are drawn from different source populations, multiplied by the between-population ePMR for *i* and *j*.

Accordingly, the ePMR is calculated as follows:

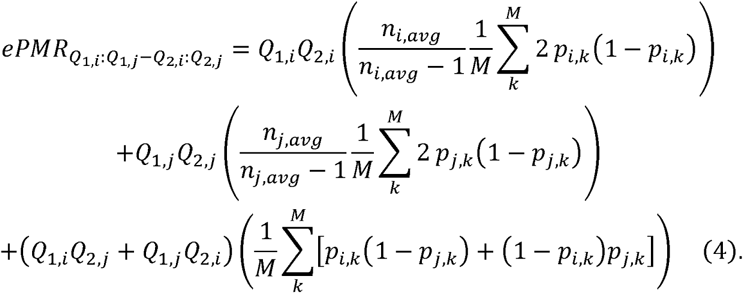

Here *Q*_1,*i*_ and *Q*_1,*j*_ are the ancestry proportions of the individual 1 from populations *i* and *j*, respectively, and *Q*_2,*i*_ and *Q*_2,*j*_ are corresponding ancestry proportions for the individual 2. We apply the finite-sample correction to each within-population diversity estimate. The between-source mismatch does not require this correction since the sources are sampled independently.

*mixKIN* then compares oPMR with this pair-specific ePMR to estimate kinship. The framework can be extended to additional source populations by summing all within- and between-source combinations. For a single homogeneous population, it retains the corrected within-population mismatch baseline described above.

### Classification Boundaries to estimate the degrees of relatedness

For non-inbred pedigree relationships, the expected kinship coefficient (*θ*) decays exponentially with the degree of relatedness, according to 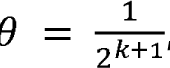, where *k* is the degree of relatedness (e.g. a value of k=0 corresponds to monozygotic twins, k=1 to first-degree relatedness, etc.)

For real data, theta can take any decimal value, and we need to define thresholds to infer discrete degrees or relatedness. In our implementation, given theta is not a linear function of k, we define classification thresholds at fractional degrees (the half degrees of relatedness) rather than using arithmetic midpoints as is done in *READv2*. Accordingly, the implemented thresholds are 0.3535 (*k*=0.5) between monozygotic twins and first-degree relatives, 0.1767 (*k*=1.5) between first- and second-degree relatives and 0.0883 (*k*=2.5) between second-and third-degree relatives at.

For unrelated pairs, the expected kinship coefficient is zero, corresponding to a degree of relatedness (k) that tends to infinity. A geometric midpoint on the degree scale between third-degree and unrelated pairs would therefore also tend to infinity, yielding a coefficient threshold of zero. We instead used the arithmetic midpoint of 0.03125 between third-degree and unrelated individuals.

### Software Implementation

*mixKIN* is implemented as a Python package accepting both EIGENSTRAT (Patterson et al., 2006, 2012) and PLINK binary (.bed, .bim, .fam) (Purcell et al., 2007) file formats as an input.

Ancestry-aware analysis requires individual ancestry proportions and source-population allele frequencies. When source populations are included in the input dataset, *mixKIN* calculates source allele frequencies and mean observed allele counts internally using supplied ploidy (i.e 2 for diploid, 1 for pseudohaploid). Alternatively, source population allele frequencies and sample sizes can be provided as input. Before calculating oPMR, *mixKIN* restricts target SNPs to those present in the source populations used to estimate the ePMR.

### Simulation Framework

As discussed above, we simulated seven independent pedigrees, each containing 14 individuals with the same structure, and combined them into a dataset of 98 individuals (Figure 1A). We used *ped-sim* (Caballero et al., 2019) with founders from 1000 Genomes Project (2015). We used the refined genetic map from Bherer et al. (2017) and crossover interference parameters from Campbell et al. (2015).

Our simulated pedigrees capture various familial relationships, including siblings, parent-offspring, grandparent-grandchild, and first cousins, providing a good framework for benchmarking the tested methods. When founders were sampled from two different populations, the pedigrees include individuals with nominal ancestry proportions of 50:50, 75:25, and 25:75.

We retained the same pedigree structure across five demographic scenarios:

i. <u>Distant-population admixture:</u> founders were drawn from Japanese in Tokyo (JPT) and Toscani in Italy (TSI).
ii. <u>Genetic outliers:</u> a predominantly JPT dataset comprising only JPT founders from six JPT-TSI pedigrees and one complete JPT-TSI pedigree, in which TSI founders and admixed individuals would be considered genetic outliers because their genetic profile differs from that of the predominantly JPT population.
iii. <u>Intra-population heterogeneity</u>: founders were drawn from the Peruvian (PEL) population, which have varying European and Indigenous American ancestry proportions. We selected founders whose two-way *qpAdm* (Haak et al., 2015) models using Iberian (IBS) and Mayan sources were not rejected at p > 0.05, while maximising estimated ancestry differences among founders.
iv. <u>Close-population admixture</u>: founders were drawn from TSI and Utah residents with Northern and Western European ancestry (CEU).
v. <u>Homogeneous population</u>: all founders were drawn from JPT.

We prepared founder genotypes from the 1000 Genomes VCF file phased with SHAPEIT2 (Delaneau et al., 2013). We restricted variants to autosomal, biallelic SNPs in the AADR 1240k panel (Mallick et al., 2024). Filtering with BCFtools using *‘-m2 -M2 -v snps’* yielded approximately 1.1 million SNPs.

Using ped-sim, we generated pseudohaploid data across a range of coverage settings: 0.03x, 0.06x, 0.10x, 0.15x, 0.25x, 0.5x, and 1x using ‘*--pseudo_hap COVERAGE*’ parameters. Coverage levels simulated independently, rather than obtained by downsampling. Finally, we converted the resulting VCF files to PLINK binary format (.bed, .bim, .fam) using *PLINK2* (Chang et al., 2015)

## Ancestry estimation and reference configurations

We estimated individual ancestry proportions for *mixKIN* using *qpAdm* (Haak et al., 2015) in *ADMIXTOOLS 2* (Maier et al., 2023). The left and right populations are provided in Supplementary Note 1, and estimated proportions are reported in Supplementary Tables S1-S5. For the homogenous JPT scenario, only one source population was used, so *mixKIN* estimated the mismatch baseline without an admixture model (*Q_JPT_* = 1).

*mixKIN* calculated source allele frequencies and mean observed allele counts internally using JPT and TSI for the distant-admixture and genetic-outlier scenarios; IBS and Mayan for PEL; TSI and CEU for close-admixture and JPT for the homogeneous scenario. We also evaluated the distant-admixture scenario using two-diploid Han individuals and two diploid French individuals from HGDP (Bergström et al., 2020) as proxies for JPT and TSI, respectively.

We used unsupervised *ADMIXTURE* (Alexander et al., 2009) to estimate ancestry proportions and source allele frequencies for *RelateAdmix* across all scenarios following (https://www.popgen.dk/software/index.php/RelateAdmix). In the distant-admixture scenario, we also tested *mixKIN* using ancestry proportions and source allele frequencies estimated by supervised and unsupervised *ADMIXTURE*. For supervised *ADMIXTURE*, we used 20 diploid individuals from each of JPT and TSI populations. For unsupervised *ADMIXTURE*, we selected individuals with an ancestry proportion > 0.99 for each cluster. We then approximated each cluster’s mean observed allele count by multiplying the number of selected individuals by their mean genotype call rate adjusted 1 for pseudohaploid, 2 for diploid data. This count was used for the finite-sample correction.

### Benchmarking

We compared *mixKIN* with *correctKin* (Nyerki et al., 2023) and *RelateAdmix* (Moltke & Albrechtsen, 2014), which account for population structure and admixture. We also included *READv2* (Alaçamlı et al., 2024) representing a commonly used approach that assumes a homogeneous population background.

#### mixKIN

We applied *mixKIN* to PLINK input files using the ancestry and reference configurations described above.

#### correctKin

We followed the analysis described by Nyerki et al. (2023), merging our simulated datasets with external reference panels of present-day individuals sourced from the 1240K AADR dataset (Mallick et al., 2024). The panels included 1,247 Eurasian individuals for the distant population admixture (JPT-TSI) and genetic outlier scenarios; 647 West Eurasian individuals for the close population admixture (TSI-CEU); 538 East Eurasian individuals for the homogeneous population (JPT); and 707 West Eurasian and Indigenous American individuals for the intra-population heterogeneity (PEL) (See Supplementary Note 2 for the reference populations).

We pseudo-haploidized all individuals using the *pseudoHaploidize* from the *correctKin* framework, followed by analysis with *PCAngsd* (Meisner & Albrechtsen, 2018). We then used the *markerOverlap* function from the *correctKin* to calculate the pairwise SNP-overlap fractions. We then used *filterRelates with these fractions and the PCAngsd* kinship matrix (.kinship.npy) to correct for SNP overlap and classify relationships. Following Lefeuvre et al. (2026), we recorded pairs classified as “uncertain” as unrelated. We also recorded relationships more distant than third-degree as unrelated. *PCAngsd*’s kinship function was originally an implementation of the *PC-Relate* algorithm, which uses principal component analysis (Conomos et al., 2016). However, the kinship estimation feature was removed in subsequent *PCAngsd* releases (post-v0.99), with developers noting it was a prototype that had not been fully tested (https://github.com/Rosemeis/pcangsd/issues/72).

#### RelateAdmix

We applied *RelateAdmix* using ADMIXTURE-derived ancestry populations and source allele frequencies, following the official documentation (https://www.popgen.dk/software/index.php/RelateAdmix). *RelateAdmix* outputs Cotterman coefficients *k*_0_,*k*_1_,*k*_2_ (Cotterman, 1940) rather than a direct degree of relatedness. We converted these outputs into a kinship coefficient (*θ*) using the standard formula: θ = 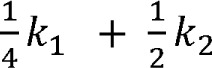. We then classified pairs using the same classification boundaries as for *mixKIN* (see above).

#### READv2

Since both *READv2* (Alaçamlı et al., 2024) and *mixKIN* calculate pairwise mismatch ratios (PMR) genome-wide, the underlying PMR estimates are mathematically identical for any given set of overlapping SNPs. Therefore, we used our own PMR estimates and normalized them using the dataset-wide median, replicating *READv2*’s default approach to calculate the kinship coefficient (*θ*) and assign degrees of relatedness based on their defined arithmetic midpoint thresholds. We omitted its minimum overlap restriction for third-degree relationships, to ensure a direct comparability with *mixKIN*.

#### Empirical application (Eastry Updown case)

We first modeled the ancestry of EAS003 and the previously inferred relatives EAS001, EAS002, and EAS006 from the medieval Eastry Updown cemetery (Sayer et al., 2025) using qpAdm with British (GBR), French, and Yoruba (YRI) populations (see Supplementary Note 1; Supplementary Table 5). We then applied *mixKIN* and compared the resulting relationship classifications with those reported in Sayer et al. (see Supplementary Note 3; Supplementary Table S11).

### Performance Evaluation Uniform Ordinal Classification

We used the 1 - UOC (Uniform Ordinal Classification) index (Silva et al., 2018). Because relatedness estimation is an inherently ordinal problem, metrics that treat all misclassifications equally are insufficient. The UOC index addresses this limitation by applying a distance-aware penalty, where the magnitude of the penalty scales with the severity of the error (Silva et al., 2018). For example, incorrectly classifying a first-degree pair as unrelated is penalized more heavily than a one-step misclassification from first-degree to second-degree. The UOC index is calculated as:

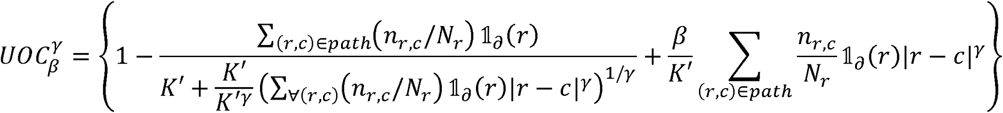

where *r* and *c* denote the row and column indices of the confusion matrix, *n_r_*_,*c*_ is the observation count in cell (*r*, *c*), and *N_r_* represents the total observations in row *r*. The term 1*_∂_*(*r*) is an indicator function that equals 1 if row r contains observations (and 0 otherwise), while *K*′ is the total number of observed classes. The parameters and control the baseline penalty for deviating from the diagonal and its distance-based scaling, respectively. Following Silva et al. (2018) and its subsequent implementation by Lefeuvre et al. (2026), we resolved these parameters by setting *γ* = 1 and integrating the metric over the range *β* ∈ [0,1]. The final index is then calculated by minimizing this cost across all consistent paths across the matrix. We integrated the R implementation of this metric from the BADGER pipeline (Lefeuvre et al., 2026) to systematically benchmark our results.

### Root Mean Square Deviation (RMSD)

To quantify the magnitude of error in estimated kinship coefficients, we calculated the root mean square deviation (RMSD) across N evaluated pairs as follows:

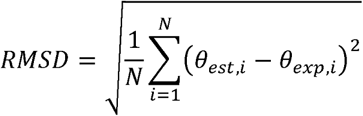

Here, *θ_exp_* and *θ_est_* are the expected and estimated coefficients for pair i, respectively. Lower RMSD indicates smaller deviations from the specified expectations.

### Mean Bias Error (MBE)

To measure the bias in the estimates, we calculated the mean bias error (MBE) as the average difference between estimated and expected kinship coefficients:

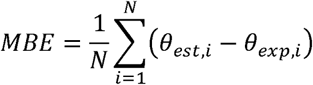

Positive values indicate average overestimation, and negative values indicate average underestimations. Values near zero indicate little overall bias towards over- or underestimation, but do not necessarily imply all estimates are accurate for individual pairs, because opposing errors can cancel each other.

## Availability of data and materials

*mixKIN* is open-source software, available on GitHub (github.com/mskilic/mixKIN).

The analysis used publicly available data from the Allen Ancient DNA Resource (AADR v62.0) (Mallick et al., 2024), available at doi.org/10.7910/DVN/FFIDCW (accessed 17 September 2025), and the 1000 Genomes Project (1000 Genomes Project Consortium et al., 2015), available at ftp.1000genomes.ebi.ac.uk/vol1/ftp/release/20130502/ (accessed 26 September 2025).

## Acknowledgements

We thank Mehmet Somel for helpful discussion and Vagif Mammedzada for testing the software, and all members of the Evolutionary Genomics Group (EGG) at UNIL for their support, suggestions, and comments. The work was funded by a Swiss National Science Foundation (PP00P3_176977) grant to A.-S.M.

