## Supplementary Material for "mixKIN: An ancestry-aware kinship estimation method for low coverage genomes"

This file contains Figures S1-S5 and Supplementary Notes 1-3, which support the methodology and main results.

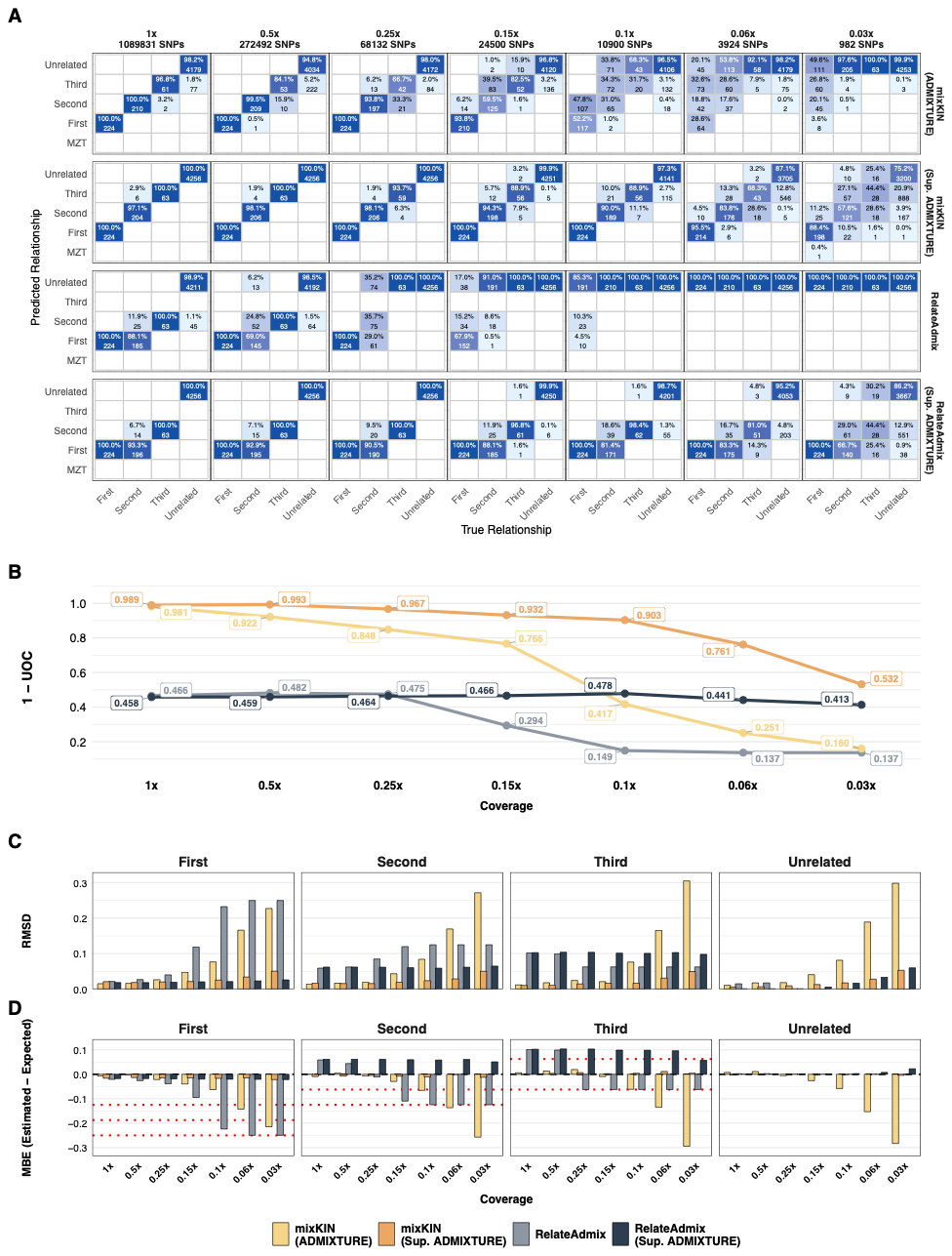

**Supplementary Figure 1: Comparison of mixKIN and RelateAdmix using ADMIXTURE-derived inputs in the distant-population admixture scenario (JPT-TSI).** The plot illustrates the accuracy of both tools when individual ancestry proportions and source population allele frequencies are estimated using either unsupervised *ADMIXTURE* or supervised *ADMIXTURE* (Sup. ADMIXTURE), demonstrating the downstream impact of upstream estimate distortions at ultra-low coverages. A reference set of n=20 diploid individuals each from the TSI and JPT populations was used for the supervised analysis. **(A)** Confusion matrices comparing true and predicted relationships for each method. Each cell shows the number and percentage of pairs assigned to corresponding predicted relationship categories. Diagonal cells represent correct classifications. Darker blue shading indicates higher values across all cells. The average number of overlapping SNPs is shown for each coverage level on top of each column. **(B)** Classification performance measured using the 1-UOC index across 1x to 0.03x coverage, higher values indicate better performance. *mixKIN* is compared with *RelateAdmix* **(C)** Root Mean Square Deviation (RMSD) between estimated and expected kinship coefficients, stratified by true relationship degree. **(D)** Mean Bias Error (MBE) calculated as $\theta_{est}-\theta_{exp}$, within each true relationship category. Positive and negative values indicate overestimation and underestimation, respectively. Red dotted lines indicate deviations corresponding to one degree of relatedness in either direction.

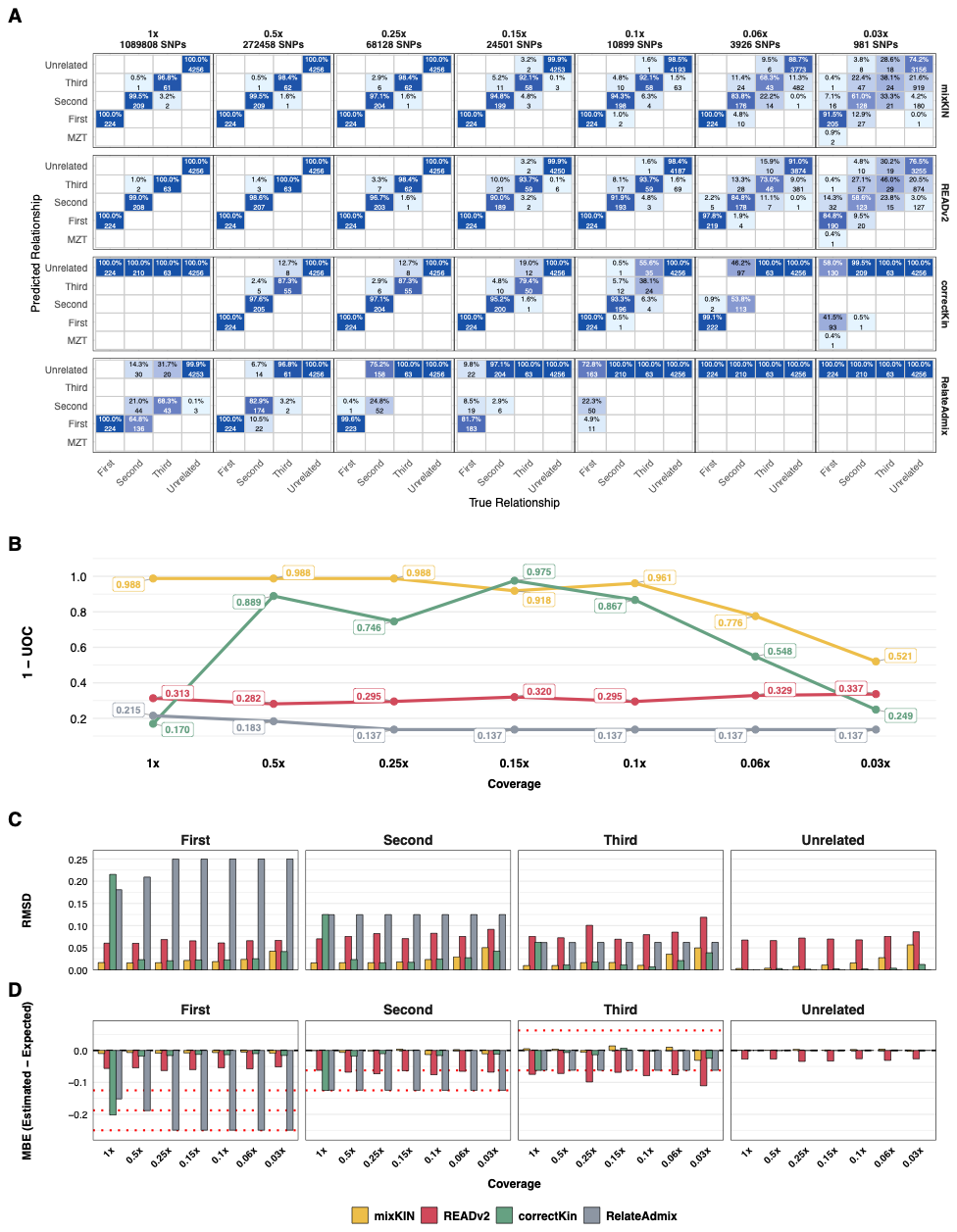

**Supplementary Figure 2: Comparison of kinship estimation methods in the genetic outliers scenario. (A)** Confusion matrices comparing true and predicted relationships for each method. Each cell shows the number and percentage of pairs assigned to corresponding predicted relationship categories. Text within each cell is written in black for percentages below 50% and in white for percentages of 50% or higher. Diagonal cells represent correct classifications. Darker blue shading indicates higher values across all cells. The average number of overlapping SNPs is shown for each coverage level on top of each column. **(B)** Classification performance measured using the 1-UOC index across 1x to 0.03x coverage, higher values indicate better performance. *mixKIN* is compared with *correctKin*, *RelateAdmix* and *READv2*. **(C)** Root Mean Square Deviation (RMSD) between estimated and expected kinship coefficients, stratified by true relationship degree. **(D)** Mean Bias Error (MBE) calculated as $\theta_{est}-\theta_{exp}$, within each true relationship category. Positive and negative values indicate overestimation and underestimation, respectively. Red dotted lines indicate deviations corresponding to one degree of relatedness in either direction.

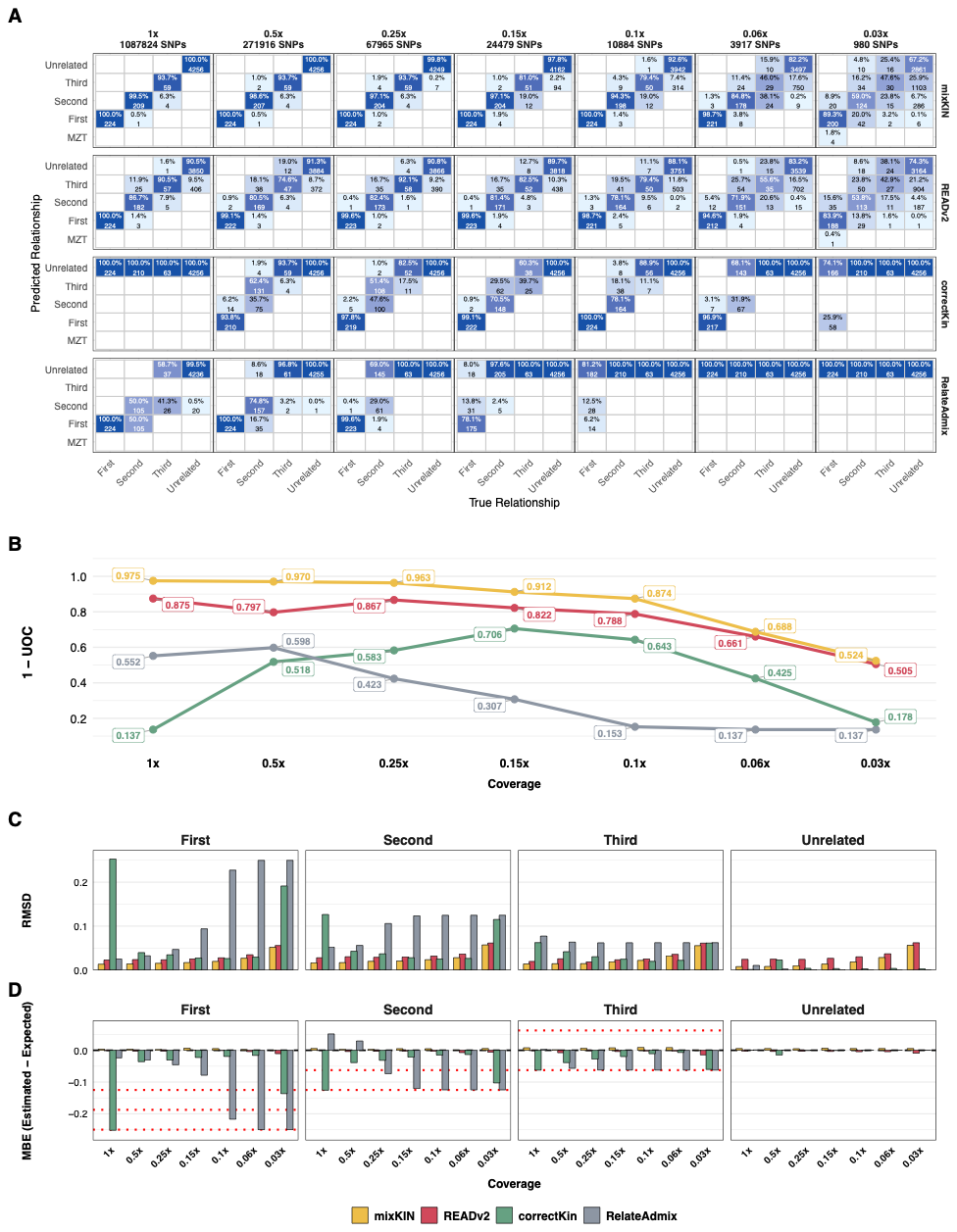

**Supplementary Figure 3: Comparison of kinship estimation methods in intra-population heterogeneity scenario (PEL).** **(A)** Confusion matrices comparing true and predicted relationships for each method. Each cell shows the number and percentage of pairs assigned to corresponding predicted relationship categories. Text within each cell is written in black for percentages below 50% and in white for percentages of 50% or higher. Diagonal cells represent correct classifications. Darker blue shading indicates higher values across all cells. The average number of overlapping SNPs is shown for each coverage level on top of each column. **(B)** Classification performance measured using the 1-UOC index across 1x to 0.03x coverage, higher values indicate better performance. *mixKIN* is compared with *correctKin*, *RelateAdmix* and *READv2*. **(C)** Root Mean Square Deviation (RMSD) between estimated and expected kinship coefficients, stratified by true relationship degree. **(D)** Mean Bias Error (MBE) calculated as $\theta_{est}-\theta_{exp}$, within each true relationship category. Positive and negative values indicate overestimation and underestimation, respectively. Red dotted lines indicate deviations corresponding to one degree of relatedness in either direction.

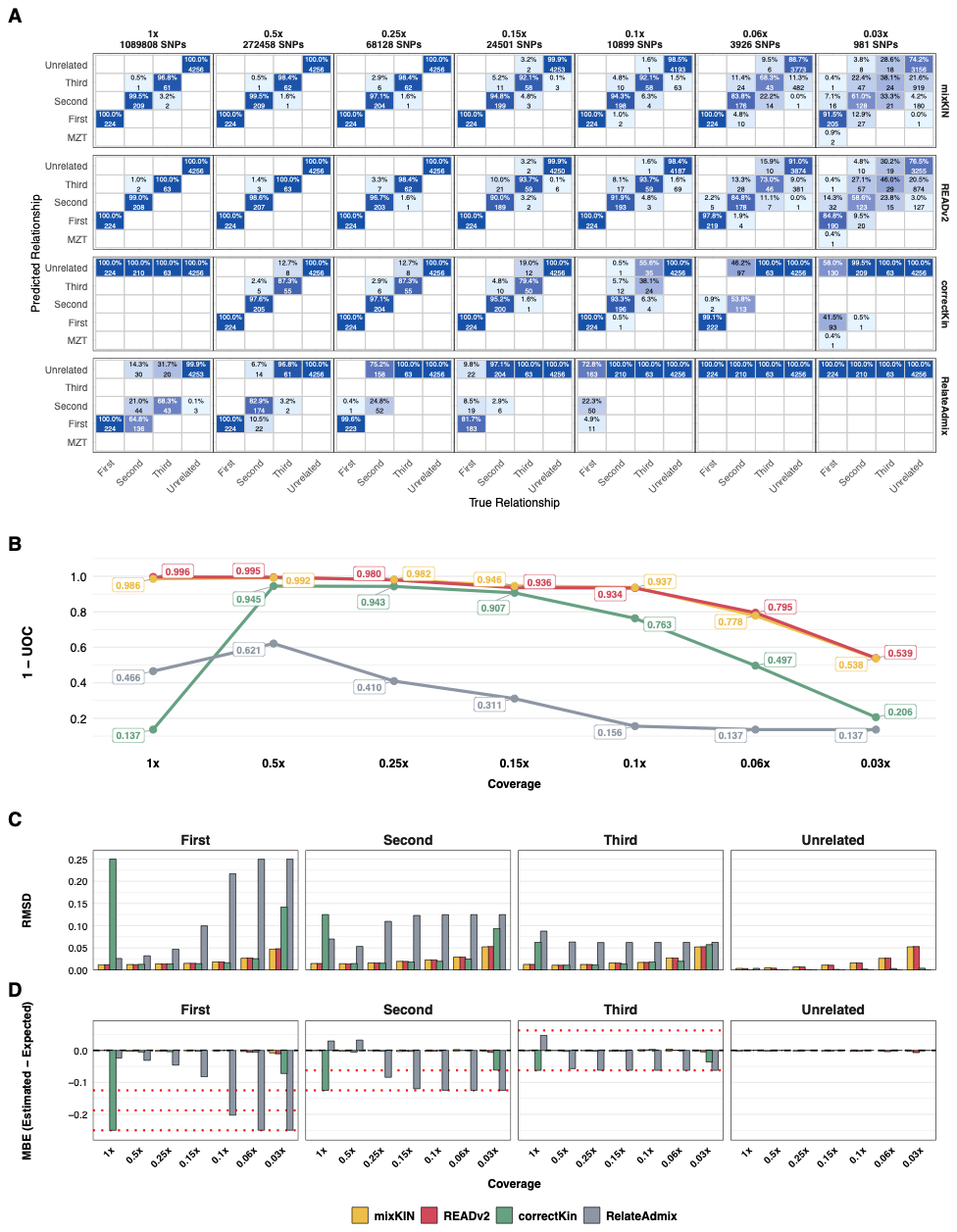

**Supplementary Figure 4: Comparison of kinship estimation methods in the close-population admixture scenario (TSI-CEU).** **(A)** Confusion matrices comparing true and predicted relationships for each method. Each cell shows the number and percentage of pairs assigned to corresponding predicted relationship categories. Text within each cell is written in black for percentages below 50% and in white for percentages of 50% or higher. Diagonal cells represent correct classifications. Darker blue shading indicates higher values across all cells. The average number of overlapping SNPs is shown for each coverage level on top of each column. **(B)** Classification performance measured using the 1-UOC index across 1x to 0.03x coverage, higher values indicate better performance. *mixKIN* is compared with *correctKin*, *RelateAdmix* and *READv2*. **(C)** Root Mean Square Deviation (RMSD) between estimated and expected kinship coefficients, stratified by true relationship degree. **(D)** Mean Bias Error (MBE) calculated as $\theta_{est}-\theta_{exp}$, within each true relationship category. Positive and negative values indicate overestimation and underestimation, respectively. Red dotted lines indicate deviations corresponding to one degree of relatedness in either direction.

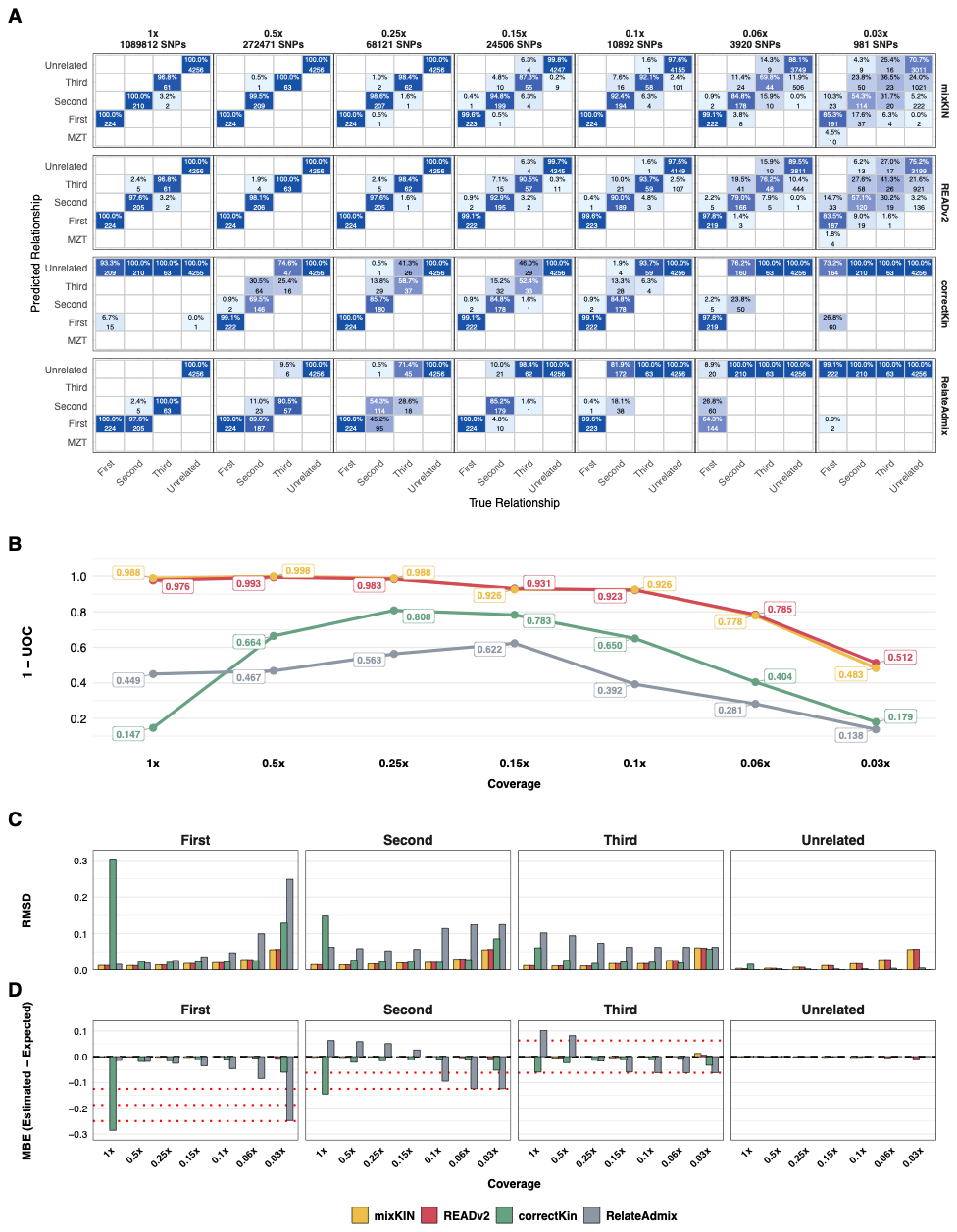

**Supplementary Figure 5: Comparison of kinship estimation methods in the homogeneous population scenario (JPT).** **(A)** Confusion matrices comparing true and predicted relationships for each method. Each cell shows the number and percentage of pairs assigned to corresponding predicted relationship categories. Text within each cell is written in black for percentages below 50% and in white for percentages of 50% or higher. Diagonal cells represent correct classifications. Darker blue shading indicates higher values across all cells. The average number of overlapping SNPs is shown for each coverage level on top of each column. **(B)** Classification performance measured using the 1-UOC index across 1x to 0.03x coverage, higher values indicate better performance. *mixKIN* is compared with *correctKin*, *RelateAdmix* and *READv2*. **(C)** Root Mean Square Deviation (RMSD) between estimated and expected kinship coefficients, stratified by true relationship degree. **(D)** Mean Bias Error (MBE) calculated as $\theta_{est}-\theta_{exp}$, within each true relationship category. Positive and negative values indicate overestimation and underestimation, respectively. Red dotted lines indicate deviations corresponding to one degree of relatedness in either direction.

**Supplementary Note 1: qpAdm Population Models**

To estimate individual ancestry proportions for the simulated demographic scenarios and the empirical case, we used qpAdm. The target source populations (left populations) and outgroup reference populations (right populations) used for each model are detailed below. The sample sizes for the left populations in these models are diploid and match the exact sample sizes used to calculate source allele frequencies in the mixKIN algorithm.

**1. Distant Population Admixture (JPT - TSI)**

**Left Populations (Exact Model):** JPT.DG, TSI.DG ($n=20, 20$)

**Left Populations (Surrogate Model):** Han.DG, French.DG ($n=2, 2$)

**Right Populations:** YRI.DG, Adygei.DG, Yakut.DG, Karitiana.DG, Russia_Kostenki14_UP.SG, Pathan.DG, Russia_MA1_UP.SG

**2. Genetic Outliers (JPT - TSI)**

**Left Populations:** JPT.DG, TSI.DG ($n=2, 2$)

**Right Populations:** YRI.DG, Adygei.DG, Yakut.DG, Karitiana.DG, Russia_Kostenki14_UP.SG, Pathan.DG, Russia_MA1_UP.SG

**3. Close Population Admixture (TSI - CEU)**

**Left Populations:** TSI.DG, CEU.DG ($n=2, 2$)

**Right Populations:** YRI.DG, Adygei.DG, Turkey_Marmara_Barcin_N.AG, Russia_Samara_EBA_Yamnaya.AG, Yakut.DG, Karitiana.DG, Russia_Kostenki14_UP.SG, Pathan.DG, Russia_MA1_UP.SG

**4. Single Heterogeneous Population (PEL)**

**Left Populations:** IBS.DG, Mayan.DG ($n=103, 18$)

**Right Populations:** YRI.DG, Adygei.DG, Yakut.DG, Karitiana.DG, Russia_Kostenki14_UP.SG, Pathan.DG, Russia_MA1_UP.SG

**5. Empirical case (Eastry Updown)**

**Left Populations:** French.DG, GBR.DG, YRI.DG ($n=20, 20, 20$)

**Right Populations:** Mbuti.DG, Ami.DG, Russia_MA1_UP.SG, Turkey_Marmara_Barcin_N.AG, Iran_GanjDareh_N.AG, Georgia_Kotias_Mesolithic.SG, Russia_Samara_HG.AG, Russia_YuzhniyOleniyOstrov_Mesolithic.AG, Serbia_IronGates_Mesolithic.AG

**Supplementary Note 2: correctKin Reference Populations**

To evaluate the performance of correctKin, which relies on PCAngsd for initial kinship estimations, we merged our simulated datasets with external reference panels of present-day diploid individuals sourced from the 1240K dataset. The specific reference populations used to capture the genetic diversity for each demographic scenario are detailed below.

**1. Distant Population Admixture (JPT - TSI)**

Abkhasian.DG, Adygei.DG, Albanian.DG, Ami.DG, Armenian.DG, Atayal.DG, Basque.DG, Bergamo.DG, Bulgarian.DG, CDX.DG, CEU.DG, CHB.DG, Chechen.DG, China_Lahu.DG, CHS.DG, Cretan.DG, Czech.DG, Dai.DG, English.DG, Estonian.DG, FIN.DG, Finnish.DG, French.DG, GBR.DG, Georgian.DG, Greek_1.DG, Greek_2.DG, Han.DG, Hungarian.DG, IBS.DG, IBS_CanaryIslands.DG, Icelandic.DG, Italian_North.DG, Italian_Sardinian.DG, Japanese.DG, JPT.DG, Korean.DG, Lezgin.DG, Miao.DG, Naxi.DG, Norwegian.DG, Orcadian.DG, Polish.DG, Russian.DG, Russia_NorthOssetian.DG, Saami.DG, Sardinian.DG, She.DG, Spanish.DG, TSI.DG, Tu.DG, Tujia.DG, Turkish.DG, Tuscan_1.DG, Yi.DG

**2. Close Population Admixture (TSI - CEU)**

Abkhasian.DG, Adygei.DG, Albanian.DG, Armenian.DG, Basque.DG, Bulgarian.DG, CEU.DG, Chechen.DG, Cretan.DG, Czech.DG, Estonian.DG, FIN.DG, French.DG, GBR.DG, Georgian.DG, Greek_1.DG, Greek_2.DG, Hungarian.DG, IBS.DG, IBS_CanaryIslands.DG, Icelandic.DG, Italian_North.DG, Italian_Sardinian.DG, Lezgin.DG, Norwegian.DG, Orcadian.DG, Polish.DG, Russian.DG, Russia_NorthOssetian.DG, Saami.DG, Sardinian.DG, TSI.DG, Turkish.DG

**3. Single Homogeneous Population (JPT)**

Ami.DG, Atayal.DG, CDX.DG, CHB.DG, China_Lahu.DG, CHS.DG, Dai.DG, Han.DG, Japanese.DG, JPT.DG, Korean.DG, Miao.DG, Naxi.DG, She.DG, Tu.DG, Tujia.DG, Yi.DG

**4. Single Heterogeneous Population (PEL)**

Abkhasian.DG, Adygei.DG, Albanian.DG, Armenian.DG, Basque.DG, Bulgarian.DG, CEU.DG, Chane.DG, Chechen.DG, Chipewyan.DG, Cretan.DG, Czech.DG, Estonian.DG, FIN.DG, French.DG, GBR.DG, Georgian.DG, Greek_1.DG, Greek_2.DG, Hungarian.DG, IBS.DG, IBS_CanaryIslands.DG, Icelandic.DG, Italian_North.DG, Italian_Sardinian.DG, Karitiana.DG, Lezgin.DG, Mayan.DG, Mixe.DG, Mixtec_1.DG, Mixtec_2.DG, Nahua.DG, Norwegian.DG, Orcadian.DG, Piapoco.DG, Polish.DG, Quechua.DG, Russian.DG, Russia_NorthOssetian.DG, Saami.DG, Sardinian.DG, Surui.DG, TSI.DG, Turkish.DG

**Supplementary Note 3: Unadjusted kinship coefficients for empirical case**

The unadjusted kinship coefficients presented in the main text were derived directly from the pairwise mismatch ratios (PMR) and the 0.255 population-average PMR baseline reported by Sayer et al. (2025), while the adjusted kinship coefficients used an individual-wise PMR baseline of 0.275 for any pair containing the admixed individual EAS003 (Supplementary Table S9).
